# A fungi hotspot deep down the ocean: explaining the presence of *Gjaerumia minor* in Equatorial Pacific bathypelagic waters

**DOI:** 10.1101/2024.01.25.577184

**Authors:** Massimo C. Pernice, Irene Forn, Ramiro Logares, Ramon Massana

## Abstract

A plant parasite associated with the white haze disease in apples, the Basidiomycota *Gjaerumia minor,* has been found in most samples of the global bathypelagic ocean. An analysis of environmental 18S rDNA sequences on 12 vertical profiles of the Malaspina 2010 expedition shows that the relative abundance of this cultured species actually increases with depth while its distribution is remarkably different between the deep waters of the Pacific and Atlantic oceans, being present in higher concentrations in the former. This is evident from sequence analysis and a microscopic survey with a species-specific newly designed TSA-FISH probe. Several hints point to the hypothesis that *G. minor* is transported to the deep ocean attached to particles, and the absence of *G. minor* in bathypelagic Atlantic waters could then be explained by the absence of this organism in surface waters of the equatorial Atlantic. The good correlation of *G. minor* biomass with recalcitrant carbon and free-living prokaryotic biomass in South Pacific waters, together with the identification of the observed cells as yeast and not as a resting spore (teliospore), point to the possibility that once arrived at deep layer this species keeps on growing and thriving.

## Introduction

The bathypelagic ocean is one of the largest reservoirs of carbon on the planet, both particulate and dissolved. The bioavailable part of this deep carbon pool, also known as labile carbon, decreases along the conveyor belt, following the aging of the water masses, being higher in the Atlantic Ocean compared to the Pacific Ocean^1^. Despite this general decrease, fast-sinking episodes, such as the collapse of a bloom, could inject fresh organic carbon into the deep sea^2^. The remanent part, known as refractory carbon, has been operationally defined as the organic carbon that cannot be degraded at in situ conditions^3^ and increases towards Pacific waters. This increase of refractory carbon along the conveyor belt is mainly explained by the fact that microbial communities act on the total bathypelagic carbon pool reshaping its composition. Among these active microbes, fungi have been proposed to play an important role as decomposers of organic matter, both labile and refractory^4^, with a specialized skillset for the second pool. Despite their importance for the microbial carbon pump, the study of bathypelagic fungal community and marine fungi in general has long been overlooked.

Pelagic fungi, which are consistently detected in seawater samples, belong mainly to two taxonomic divisions, Ascomycota and Basidiomycota^5^. Species affiliated with these groups, including *G. minor*, survive thanks to two main trophic strategies: saprotrophy and parasitism. Saprotrophic fungi degrade and recycle organic matter with extracellular enzymes^6^, in particular, it has been reported that the abundance of CAZymes (Carbohydrate Active enZymes) increases toward mesopelagic waters^7^ pointing to the central role of fungal catabolism in aphotic environments. A similar pattern was also detected for genes related to protein degradation^8^. The utilization of extracellular enzymes is a more effective digestion strategy if associated with a particulate-attached lifestyle and, actually, in bathypelagic waters fungi are main colonizers of marine snow, exceeding in some cases the biomass of bacteria^9^. As parasites, pelagic fungi are associated with different hosts, including animals, macroalgae, and microplankton^10–12^. Several environmental variables have been proposed to shape fungi abundance in photic waters, including temperature, particulate organic matter^13^, salinity, depth, oxygen and nitrate^14,15^ whereas less information is available about the drivers of their distribution in the bathypelagic realm. It has been suggested that they increase in diversity with depth, mirroring what happens with prokaryotes^5^.

During the Malaspina 2010 expedition, Basidiomycota were found to be one of the most important microbial eukaryotic groups in the Bathypelagic Ocean^16^. Their relative abundance, based on 454-pyrosequencing, peaked at equatorial Pacific waters, being the most abundant 18S rDNA sequence virtually identical to *Gjaerumia minor*. A further Illumina sequencing^17^ confirmed that one of the ASVs (Amplicon Sequence Variants) retrieved (ASV-363) has 100% similarity with *Gjaerumia minor* (NG_063045). A *G. minor* MAG retrieved from the deep ocean featured an ITS rRNA sequence identical to a strain isolated from a pleural effusion of a child with pneumonia (KT149771). This species, previously known as *Tilletiopsis minor,* was recently renamed since the entire taxonomy of *Exobasidiomycetes* has been redefined based on their phylogenetic relations^18^. Although the most frequently reported habitat of *G. minor* is plant material (dead or alive), through propagules in the air, this yeast is capable of colonizing other niches^19^, including the deep ocean. This ability to adapt to different environments is well exemplified by the fact that this fungus has been found to be a pathogen both for plants and humans. *G. minor* is one of the causes of the “white haze” in apple trees^20^, a post-harvest disease that flourishes with humidity, cold (4°C) and low oxygen levels^19^ while in literature three cases are reported of *G. minor* as a human pathogen, finding this yeast involved in subcutaneous mycosis^21^, severe pneumonia^22^ and corneal abscess^23^. The unusual plasticity of *G. minor,* coupled with its uneven distribution in the Bathypelagic Ocean, makes this interesting species worth of deeper analyses.

Here we reanalyze published sequencing datasets from the Malaspina expedition to report the distribution of the *Gjaerumia minor* along the vertical profiles of several oceanic stations. In addition, taking advantage of the fact that a strain of *Gjaerumia minor* is commercially available, we were able to develop and test a new TSA-FISH probe with the aim of confirming the presence of this fungus in marine pelagic samples, describing its morphology, quantifying its abundance and investigating a possible role in the labile and refractory carbon degradation process. Finally, we aim to highlight which abiotic and biotic parameters may explain this organism’s uneven distribution with the final goal of better characterizing the trophic dynamics of the Bathypelagic Ocean.

## Methods

Information about the studied stations and sampling of Malaspina 2010 expedition could be found in previously published works. In particular, the methodology for bacterial abundance with flow-cytometry and prokaryotic biomass calculation is described in Pernice et al. (2015)^24^; DNA extraction, 454-pyrosequencing, and bioinformatic pipeline for samples belonging to the deep ocean are in Pernice et al. (2016)^16^; DNA extraction and Illumina sequencing for 12 vertical profiles (5 to 4000 m) are in Giner et al. (2020)^17^; Illumina sequencing from samples belonging to surface of the entire cruise are in Logares et al. (2020)^25^; and data for Illumina sequencing in bathypelagic samples are in Junger et al. (2023)^26^. All the Illumina raw data (12 vertical profiles, global surface and global bathypelagic) has been newly analyzed here with DADA2^27^ to define the distribution of the ASV of interest (363) in the different datasets. Data for fluorescence Dissolved Organic Matter (FDOM) were obtained using parallel factor analyses (PARAFAC) as described in Catalá et al. (2016)^28^ and DOM data and AOU values are from Catalá et al. (2015)^29^.

For TSA-FISH analyses, a seawater sample of 475 mL was fixed with 25 mL of 37% formaldehyde (final concentration 1.85%, at least 1h at 4°C) and filtered on board on a 0.6 µm pore size polycarbonate filter (25 mm diameter). The filters were stored at -80 °C until processing. The TSA-FISH probe (Gmin01, CGACCACCATGTGCCCTT 5’-3’) to count *Gjaerumia minor* cells at the microscope was designed based on the target sequence ASV-363 as well as sequences belonging to *G. minor* from NCBI. The probe was checked in silico against the SILVA 138 database and results to be species-specific (the closer non-target sequence had 3 mismatches). The optimization of the hybridization condition was done using a commercial strain of *G. minor* (*Tilletiopsis minor* Nyland fungal strain, JCM No. 8361). A first attempt with a standard TSA-FISH protocol, as described in Pernice et al. (2015)^24,30^ adapted from Pernthaler et al. (2001)^30^, gave poor results (less than 25% of positive cells in the culture). The addition of helpers to contiguous regions of the probe (HelperA-Gmin1 5’-GCGGGCTCGCGGCGATCAAT-3’; HelperB-Gmin1 5’-ACCAAGTTTGCCCAAGTTTT-3’) increased the results to 65% of positive cells. The use of Wheat Germ Agglutinin conjugated with Alexa Fluor 594 (Thermofisher; WGA 0.01mg mL ^−1^, 30 min at RT), a fluorescent marker staining chitin, confirmed the presence of a thick wall in cells labeled with the probe so it was decided to include an additional permeabilization step. After trying several protocols, the best results were obtained using an incubation step for 1 h at 30°C in a permeabilization buffer (pH 6.5) consisting of 1x PBS, 1% SDS, 1 mg mL^−1^ chitinase and 6 mg mL ^−1^ Glucanex (a cocktail of enzymes isolated from *Trichoderma harzianum* that contains β-glucanase, cellulase, protease and chitinase as in Priest et al. 2021)^31^. With this permeabilization step we reached more than 90% of positive cells (Fig. 1). Finally, the optimal hybridization stringency was determined by keeping the temperature constant at 35°C and varying formamide concentrations (0% – 70%) in the hybridization buffer. The optimal value was 20% formamide, the highest possible concentration before probe signal intensity decreased. We then applied the developed TSA-FISH protocol to a subset of 34 environmental samples from the deepest layer of the Malaspina 2010 expedition.

**Figure 1:**
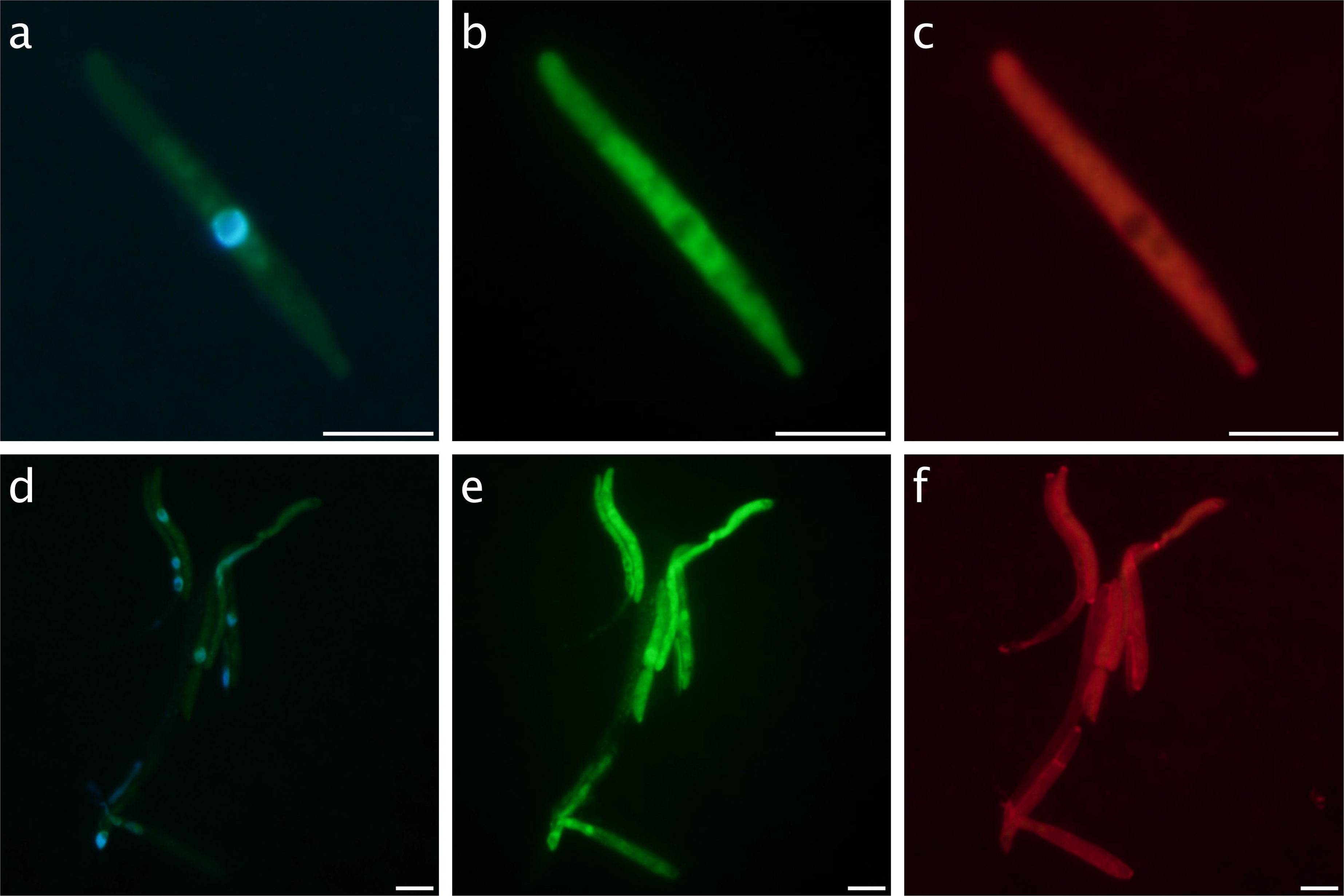
Epifluorescence pictures of cultured cells of *Gjaerumia minor* with triple staining protocols. Panels above show a single elongated cell, possible an hypha, stained with a) DAPI, b) TSA-FISH (Gmin01 probe conjugated with Alexa-488), and c) WGA. The panels below show an entire hyphal structure stained with d) DAPI, e) TSA-FISH, and f) WGA. Nuclei are visible in blue in a and d, the ribosome-containing cytoplasm in green in b and e, and the outer chitin cover in red in c and f. Scale bar is 5 µm.

Target cells (two morphotypes, rounded and elongated) were counted by inspecting a transect between 30 and 80 mm (average of 54 mm/sample equivalent to 540 fields). Microscopic analyses were performed on an Olympus BX61 epifluorescence microscope (Olympus America Inc.) at 1000× magnification under UV for DAPI, blue light for A488 (TSA-FISH) and green light for WGA. Pictures were taken on an Olympus DP72 camera connected to the microscope. Biovolume for the rounded morphotype was calculated by assuming a spherical cell based on the average radio (100 cells measured). The biovolume of the elongated morphotype was calculated assuming a prolate spheroid shape (Hillebrand et al., 1999)^32^ based on the following formula: V=pi/6*d^2^*h, where h is the largest cell dimension and d is the largest cross-section of h, average h and d are based on 40 measured cell. We then used the equation of Menden-Deuer and Lessard (2000)^33^ to convert cell biovolume to cell biomass: pgC cell^−1^ = 0.216*(Biovolume ^0.939^). Within each sample, average cell biomass times cell abundance counted by TSA-FISH for each morphotype was calculated and then summed to obtain the total biomass of the *G. minor* population.

Statistical analyses (Regression analyses and Pearson correlation) were performed with Rstudio (package Hmisc).

## Results

### Morphotypes

Two different morphotypes have been identified as *Gjaerumia minor* by the TSA-FISH probe (Fig. 2), one rounded with an average diameter size of 1.5 µm (panels a and b), and one elongated (panels c and d) with an average h (larger cell axis) of 5.2 µm and an average d (smaller cell axis) of 1.5. The rounded morphotype, which represents the majority of cells retrieved, shows a single nucleus per cell, evidenced in the figure by DAPI stain. This cell type never appears in culture and was not stained by WGA, as it is clearly visible when comparing to the elongated shape in Figs 2g, 2h and 2i. So, it was assumed that the rounded morphotype has low chitin content. Although a rounded shape is often associated in congenera species with teliospore (resting spore, ticker chitin wall, dikarya), our observations (cell size, lack of thick chitin, and the presence of only one nucleus) lead us to consider the rounded morphotype not a resting spore but an active yeast cell. The elongated morphotypes, present only in Pacific waters, are probably *hyphae*, sometimes found in chains (Figs. 2c, 2d) but more often as single units (Figs 2g, 2h and 2i) with a visible nucleus in the middle as observed also in cultured cells.

**Figure 2:**
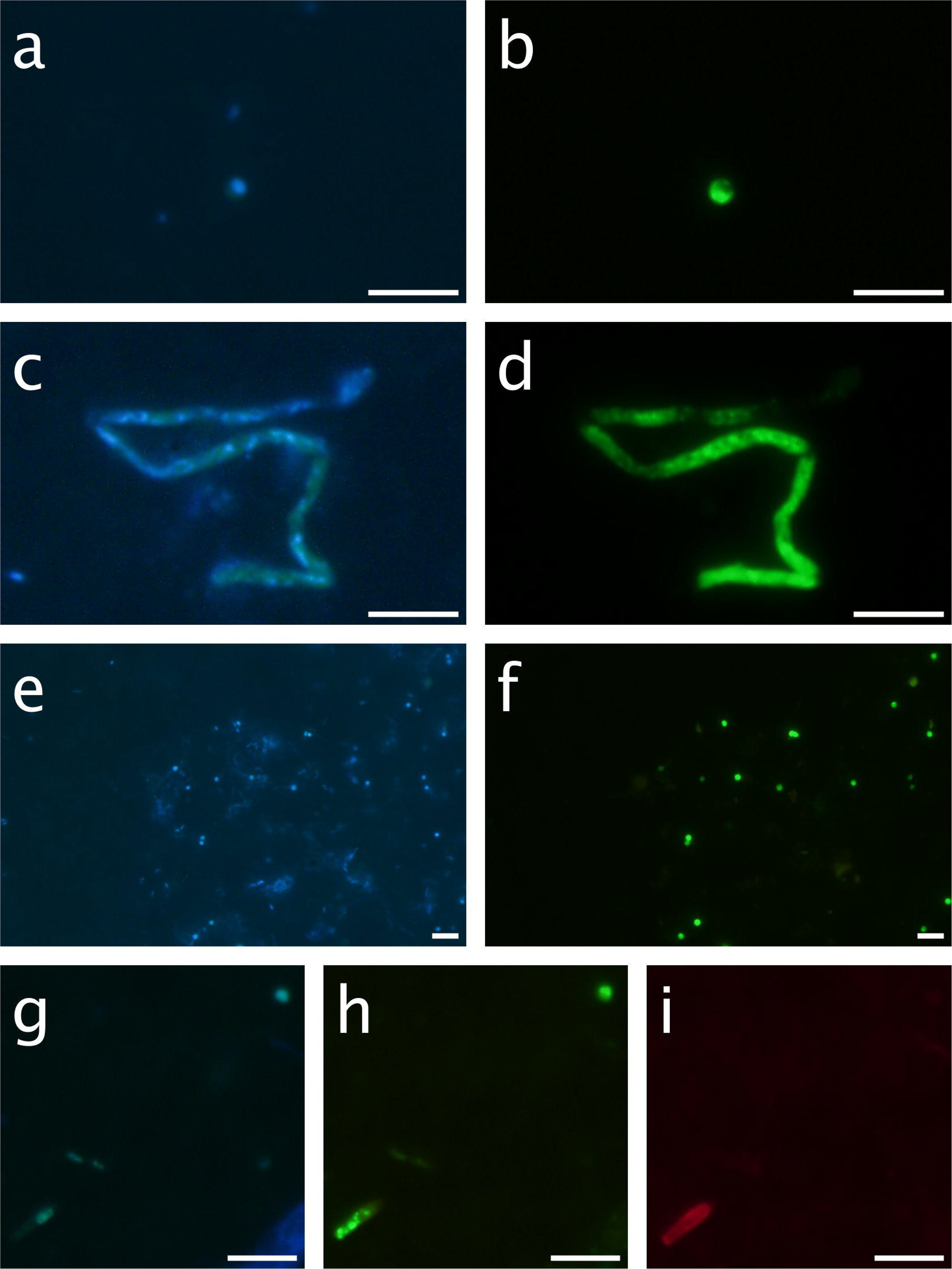
Examples of cells from environmental samples. A rounded morphotype (1.06 µm on average, probably a yeast) from ST83 stained with DAPI (a) and the Gmin01 probe (b). A hyphal structure (h= 6.9 µm, d=1.16 on average) from ST88 stained with DAPI (c) and the Gmin01 probe (d). A group of rounded morphotypes embedded in a gel particle from ST97 stained with DAPI (e) and the Gmin01 probe (f). A field including the two morphotypes from ST92, an hyphae in the low-left corner and a rounded cell in the up-right corner stained with DAPI (g), the Gmin01 probe (h) and WGA (i). Note that the chitin staining applies to the hypha but not to the rounded cell. Scale bar is 5 µm.

### Vertical distribution of the target 18S rDNA sequence

In order to have a general view of the distribution of *G. minor* in the global ocean, we used already published sequencing data of 18S rDNA genes from the 0.2-3 µm size fraction (Illumina tags grouped in ASVs). The relative abundance of the *G. minor* ASV (ASV-363) was available for 12 vertical profiles from the surface till 4,000 m and for 124 surface samples of the entire cruise (Fig. 3). *G. minor* ASV tends to increase with depth, as shown by the median of its relative abundance (excluding zero values), which is higher in the bathypelagic ocean (median of 0.02% of tags). Sequences belonging to *G. minor* were not found in 29% of the samples of the Bathypelagic realm, while this number was 53% for Mesopelagic and 67% for Epipelagic samples, further stressing its higher presence in the deeper ocean. Illumina sequencing was also performed in 22 bathypelagic samples belonging to a different size fraction (0.8-20 µm). This set of samples shows higher values with a median of 1.73% and a maximal value of 25%. The vertical distribution clearly points to the bathypelagic ocean as a preferred environment for this fungus and we decided to focus on the deepest sampling point for the TSA-FISH analysis.

**Figure 3:**
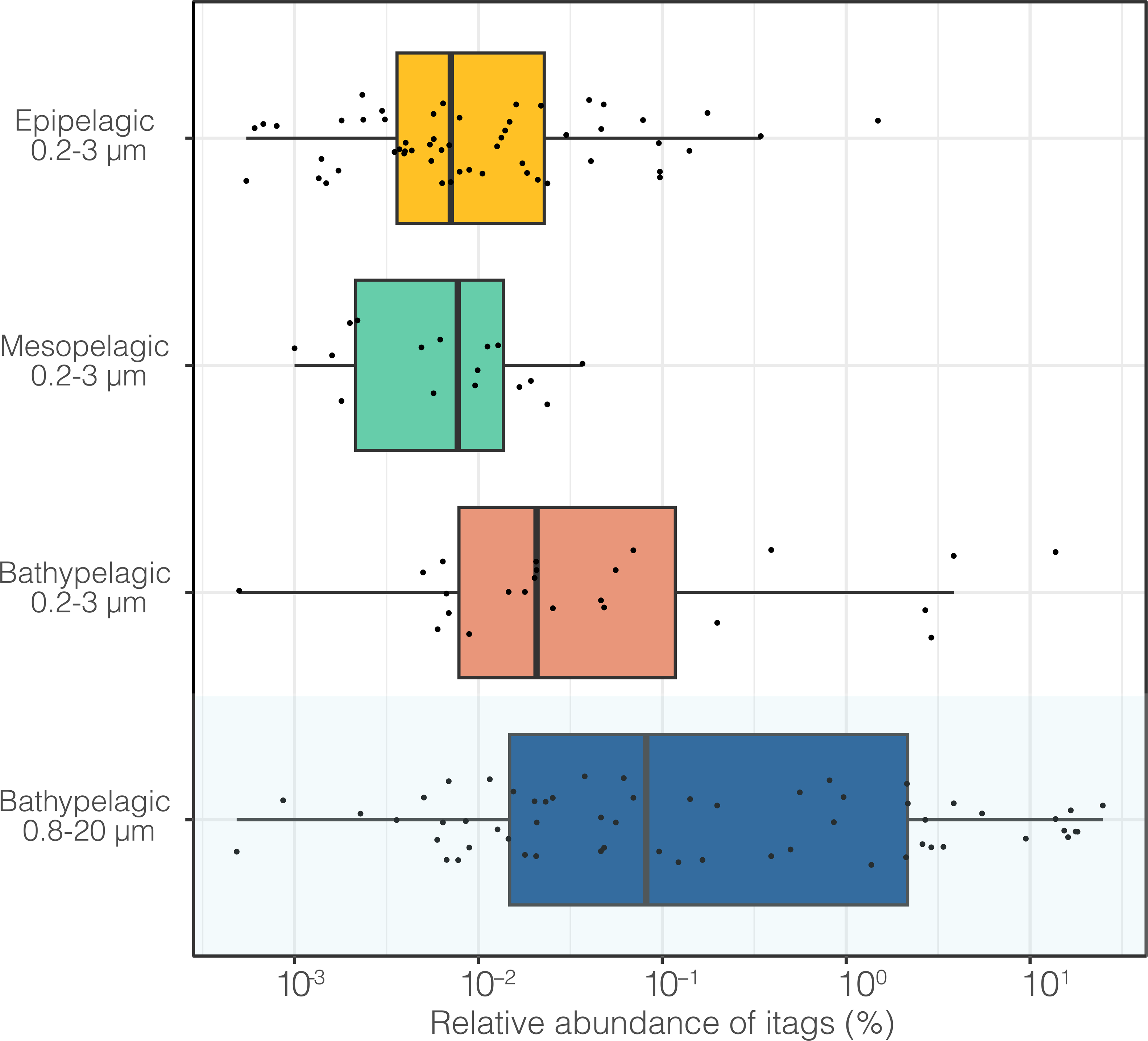
Vertical distribution of *Gjaerumia minor* based on the relative abundance of ASV-363 that is 100% similar to cultured *G. minor* (strain AB7-11). Sequences derive from previously published Illumina datasets^17, 25^. Samples from 0.2-3 µm size-fraction were grouped in three layers: Epipelagic (3-200 m), including 149 samples, Mesopelagic (200-1000 m), including 32 samples, and Bathypelagic (1000-4000), including 31 samples (zero values are not shown in the graph). The last boxplot (in blue) shows the relative abundance in 22 Bathypelagic samples belonging to the 0.8-20 µm fraction (no zero values found in this dataset).

### Cell abundance in the Bathypelagic region and fit with sequencing data

TSA-FISH analysis targeting *G. minor* was performed on the deepest sample (between 2600 and 4000 m) of 34 stations. Figure 4a shows the cell abundance of the two morphotypes retrieved along the cruise track crossing the Atlantic, Indian, and Pacific Oceans. Cells were observed in all 34 bathypelagic samples inspected, with the rounded morphotype being always more abundant than the elongated one. *G. minor* rounded cells had low abundances (less than 10 cells mL^−1^) both in Atlantic and Indian oceans whereas they were more abundant in Pacific waters, peaking at station 91 with 107 cells mL^−1^. Elongated cells were also more abundant (3-6 cells mL^−1^) where rounded cells were abundant (stations 88-91-92).

**Figure 4:**
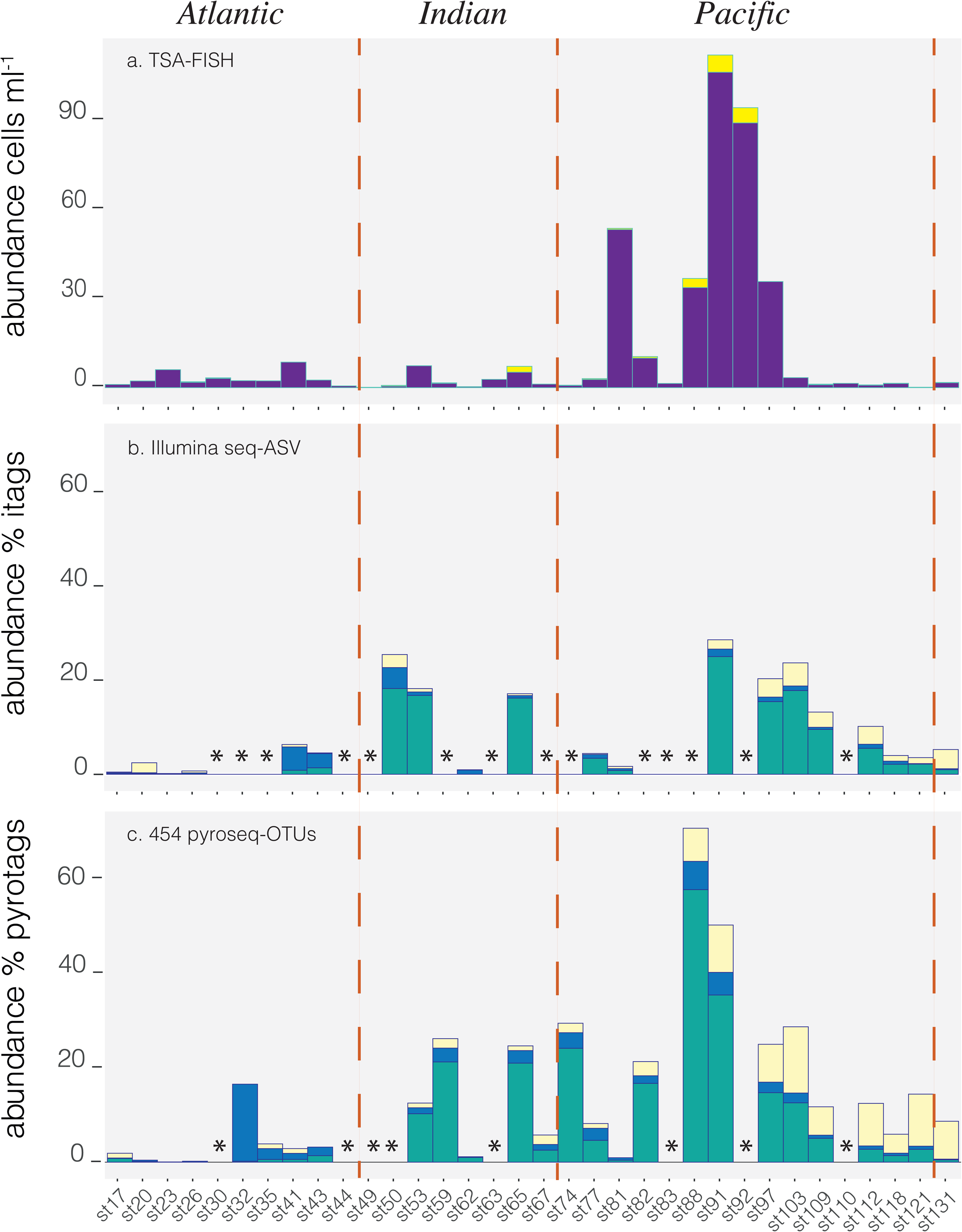
Distribution of *Gjaerumia minor* in bathypelagic waters (deepest sample in each station) along the entire track of Malaspina cruise. Oceanic boundaries are depicted as a red dotted line for the three graphs. a) Cell abundance of rounded cells (probably yeasts, violet) and elongated cells (hyphae, yellow) stained with TSA-FISH in the deepest point of 34 stations spread across Atlantic, Indian and Pacific Oceans; b) In green it is shown the relative abundance of *G. minor* illumina tags (ASV_363), abundance of tags belonging to other Basidiomycota is shown in blue whereas tags belonging to Ascomycota are shown in light yellow; c) relative abundance of 454-pyrosequenced tags (in 26 of the 34 previous stations) published in Pernice et al. (2016)^16^ (size fraction 0.8-20 µm), tags belonging to OTU-6359 (99.7 similar to *G. minor*) are shown in green, other Basidiomycota in blue and Ascomycota in yellow. Illumina and 454 sequencing were done from the same DNA extraction, asterisks show stations where no sequencing data was available.

We then compared the cell counts with the relative abundance of the corresponding phylotype in the same samples, analyzed in the 0.8-20 µm size fraction where *G. minor* had a larger representation, this fraction includes 20 samples of the Illumina survey reported before for the vertical profile (Fig. 4b) and 26 samples from a previous report^16^ in which the same DNA extracts and the same 18S DNA region were analyzed and 454-pyrosequenced (Fig. 4c). As expected, since the two sequencing analyses are based on the same DNA extract, the relative abundance of the specific OTU (by pyrosequencing) and the specific ASV (by Illumina) have a very strong direct relationship (n=19, R=0.84, p<0.001). Both sequencing datasets found an almost absence of *G. minor* in Atlantic Ocean, an intermediate presence in the Indian ocean and maximal values in the Pacific Ocean. There was a statistically significant moderate direct relationship both for the specific ASV (n=18, R=0.54, p=0.02) and the specific OTU (n=26, R=0.51, p=0.008) with the counts of rounded cells. These moderate fits could be explained by the typical errors of each approach and the huge difference in filtered volume which was ∼120 L for the DNA analysis and only 500 mL for the TSA-FISH. Sequencing data also allowed us to put the abundance of *G. minor* (in green in Figs. 4b and c) in the context of other fungi, like the rest of Basidiomycota (in blue) and Ascomycota (in light yellow). In the Indian and Pacific Oceans, where *G. minor* is abundant, this single species represents almost the totality (median of 87%) of Basidiomycota. Ascomycota is generally less abundant than Basidiomycota along the cruise track, but follow a similar global distribution, being functionally absent in the Atlantic Ocean and abundant in Pacific waters. Ascomycota maximal abundance is shifted to station 103 respect to the peak of the Basidiomycota in station 88.

### Biomass and its relation with AOU and FDOM

The biomass of the two morphotypes combined (Total Biomass) ranged between 1.74·10^−5^ and 2.05·10^−2^ µg C L^−1^, being globally the rounded morphotype the dominant contributor to the species biomass. The distribution of biomass along the entire cruise is shown in Fig. 5 (bar plot), with the highest values found in equatorial Pacific waters. The points on the graph correspond to AOU (Apparent Oxygen Utilization), which is a proxy for the age of the water mass, higher AOU values imply older waters. Considering the entire cruise track, *G.minor* biomass is higher in aged waters. A similar picture is obtained observing the relationship between *G. minor* biomass and the Fluorescent Dissolved Organic Matter (FDOM) (Fig. 6). The FDOM profile for each sampling point was obtained through a PARAFAC analysis^28^ and it is composed by four peaks, two belonging to Humic-like matter (C1 and C2) that are proxies for the refractory carbon, and two associated with protein-like matter (C3 and C4) that represent the labile carbon. Among the four FDOM components, C2 and C4 did not show a significant relationship with *G. minor* biomass nor bacteria (both in total Bathypelagic and considering only Pacific samples) and were not considered in this analysis. The correlation between fungi biomass (green dots) and FDOM other two peaks (C1-C3) as well as the relationship between bacteria biomass (blue dots) and the same parameters it is shown in Fig. 6, samples belonging to Pacific waters are in bold.

**Figure 5:**
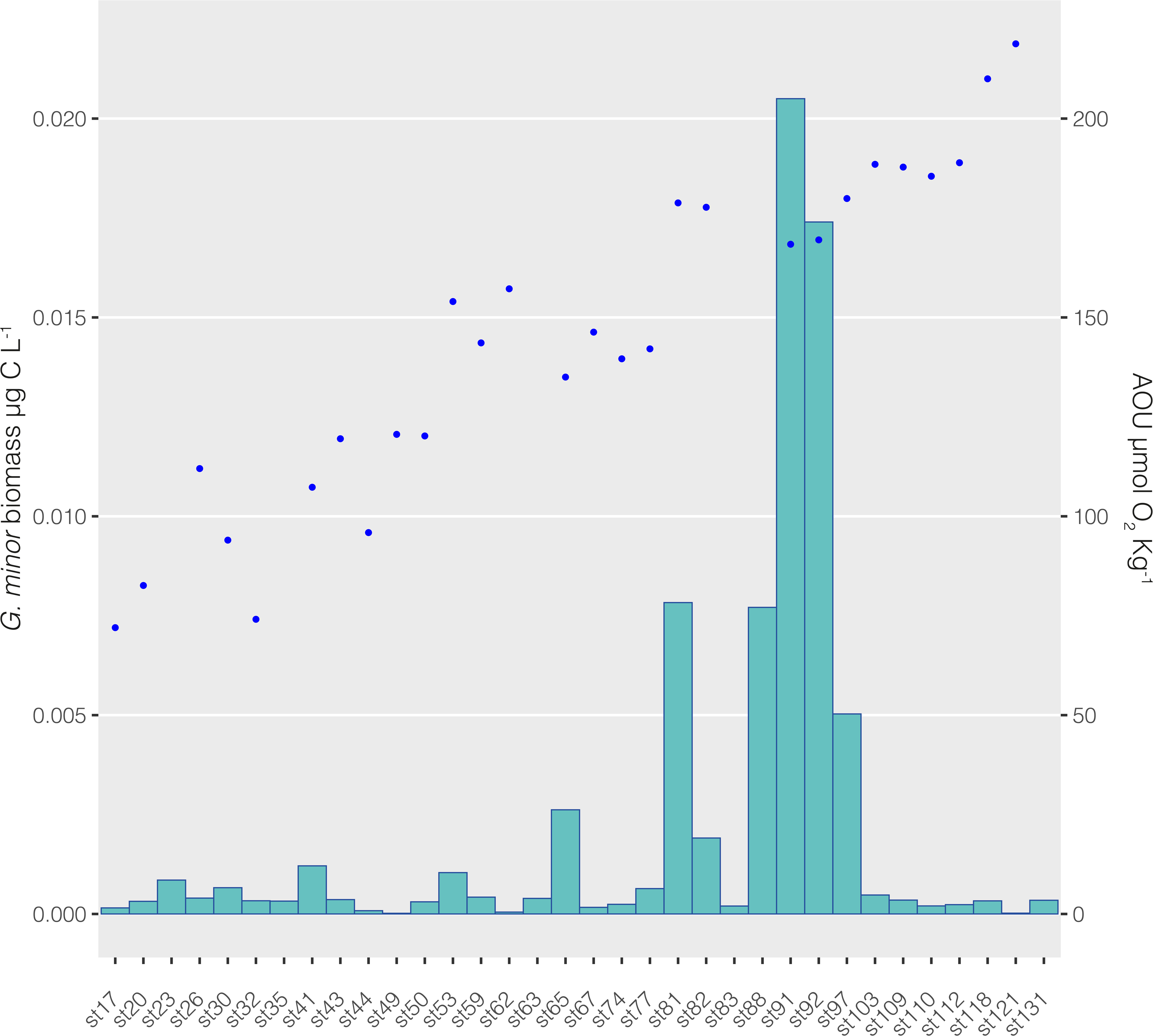
Distribution of *Gjaerumia minor* biomass and Apparent Oxygen Utilization (AOU, µmol O_2_ Kg^−1^) across the Malaspina track. The biomass (expressed as µg C l^−1^) is visualized as a barplot (left y axis) whereas the AOU, corrected by the real value of measured oxygen, is represented by the blue dot (right y axis). AOU is a measure of the amount of oxygen respired in the deep ocean, higher values of AOU are typical of aged waters.

**Figure 6:**
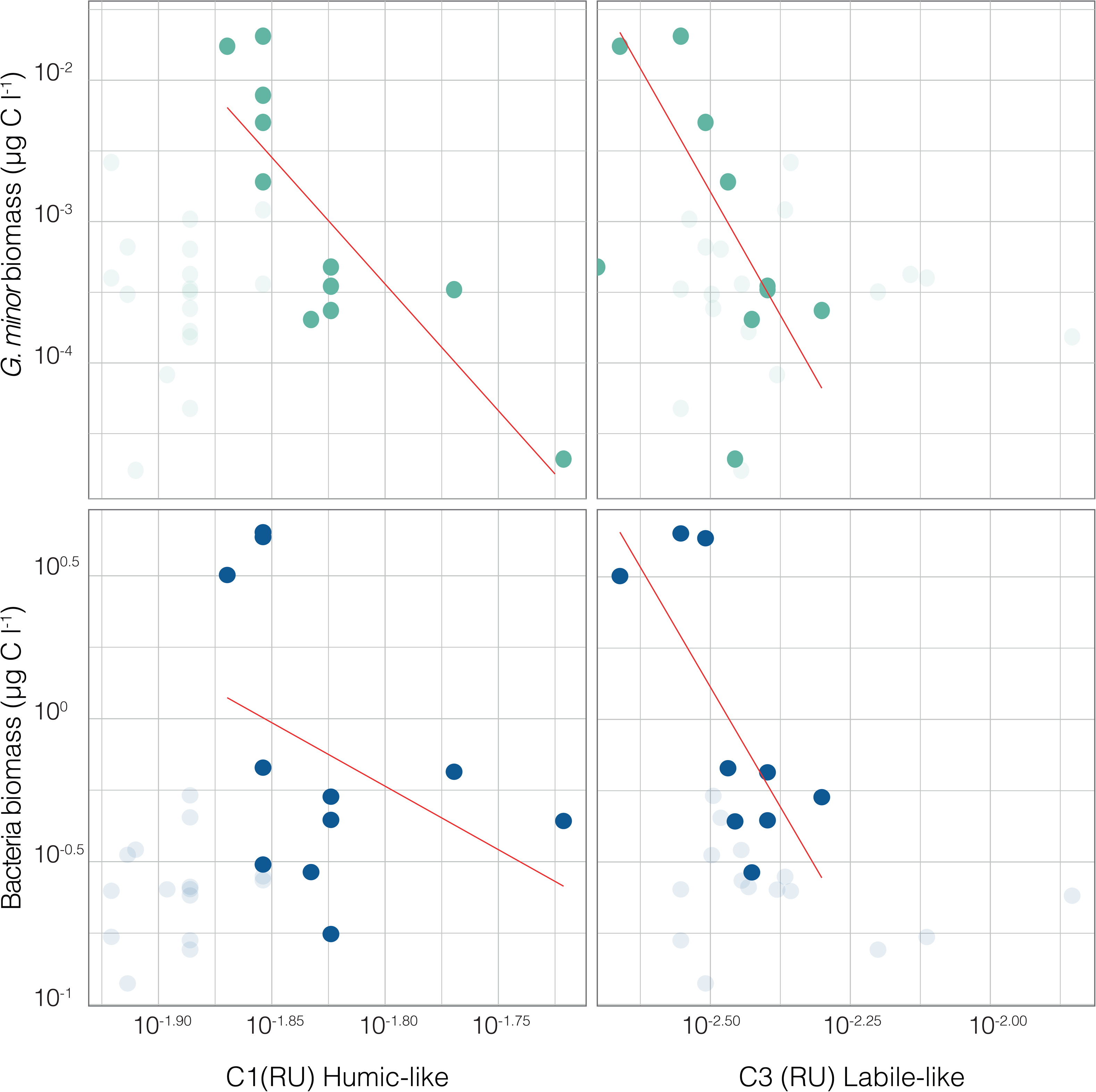
Microbial biomass relationship with C1 peak of FDOM (Humic-like, on the left) and C3 peak of FDOM (Labile-like, on the right). *G. minor* biomass is in green whereas bacterial biomass is in blue; dots belonging to Pacific samples are darker than the rest. Red lines indicate the linear relationships between biomass and FDOM values using only the darker dots.

*G. minor* in Pacific waters has a very strong inverse relationship with the Humic-like C1 peak (n=11, R=-0.83, p=0.001). So, despite the presence of *G. minor* being higher in Pacific compared to Atlantic waters, considering only the Pacific region that is richer in recalcitrant carbon (C1 peak), the abundance of this fungus is higher where the C1 concentration is lower suggesting a possible consumption of recalcitrant carbon by *G.minor*. It is important to note that, despite the biomass of bacteria is much higher than that of *G. minor*, the relationship between total bacteria biomass and C1 is not significant (p=0.22) in Pacific waters. Considering also the Pacific samples, *G. minor* biomass presents also a strong inverse relationship with the protein-like C3 peak (n=9, R=-0.72, p=0.027), which in this case is resembled by the prokaryotic biomass (n=9, R=-0.74, p=0.023).

### Abiotic and Biotic parameters

Abiotic parameters (temperature, salinity, conductivity and oxygen) did not have a significant correlation with *G. minor* biomass both for the global bathypelagic ocean and considering only Pacific samples. Higher values of biomass are within a narrow range of values both for salinity (around 34.7 psu) and oxygen (3.23-3.58 mL L^−1^), which is expected considering that they all belong to the same water mass. The values of *G.minor* biomass are extremely low compared with the total prokaryotic biomass which range between 0.12 to 4.49 µg C L^−1^. Nevertheless, the biomass of *G.minor* and prokaryotes are strongly correlated (n=26, R=0.8, p<0.001). In particular, in Pacific waters, the Low Nucleic Acid content bacteria (LNA) better correlated with the *G. minor* biomass than total bacterial biomass (R^2^=0.86 versus R^2^=0.59).

## Discussion

The presence of a clear hotspot for *G. minor* in the bathypelagic region of the Equatorial Pacific has been shown in consensus by two different sequencing analyses and a species-specific TSA-FISH probe. Since the ASV retrieved from the sequencing analyses was 100% similar to a cultured strain of *G. minor* it was possible to test the probe before its application to environmental samples. The TSA-FISH technique allowed us to visualize the morphotype (life-stage) of the targeted fungi, showing that the majority of the *G. minor* cells were rounded, with a single nucleus and unstained with WGA (Fig. 2). Larger elongated cells were also observed but at much lower abundance.

*G. minor* belongs to the class *ustilaginomycetes,* which are usually dimorphic, producing a saprobic haploid yeast phase and a parasitic dikaryotic phase^34^, so particular attention was given to the identification of the proper life-stage of the rounded cells. If they corresponded to a resting stage (teliospore) this would mean that *G. minor* is not thriving in the bathypelagic environment, whereas a yeast phase opens the possibility that *G. minor* is alive and active in those waters. Three hints suggest us to exclude the teliospore hypothesis: i) their size is too small; although teliospores from the sister species *Gjaerumia ossifragi* (Bauer 2005, figs 5-14)^35^ show a rounded shape, they are ten times larger than the ones retrieved here, and in proportion *G. minor* rounded cells seem too little for the size of the hyphae to be a teliospore, ii) they are not dikaryotic, as a teliospore is expected to be; we did not observe two nuclei in DAPI stained samples, iii) last and most relevant, rounded cells are not stained by WGA, a specific stain for chitin; teliospores, on the other hand, are expected to have a very tick chitin wall (Figs. 2 g-i). The lack of WGA staining is probably due to the fact that yeast merging and budding are favored by a thinner chitin wall. There is still the possibility that the rounded morphotype is a reproductive spore, such as basidiospore (sexual) or basidioconidia (asexual), but although the pressure of the bathypelagic environment could force a more globose shape, both of them are expected to be fusiform^35,36^, therefore, we also tend to exclude this hypothesis.

*G. minor* is known to have a strong plasticity being retrieved both as plant parasites^19,20^ and human pathogens^21–23^. In the frame of Malaspina 2010 cruise, it is often present also in surface waters (51 of 136 stations), although in a lower percentage. It is clear that *G. minor* is not an endemic species of the Bathypelagic environment but is transported there from land through the surface global ocean. For prokaryotic communities, based on samples from the same cruise, it has been reported a direct connectivity through fast-sinking particles between surface and the bathypelagic layer^37,38^. We hypothesized a similar mechanism of transport for *G. minor*, as previously suggested by Bochadansky and colleagues in 2017^9^. This idea is also reinforced by the fact that the osmotrophic food acquisition in fungi, based on the secretion of extracellular enzymes, better suits a particle-attached life-style^6^. These particles could be represented by sinking *G. minor* hosts (phytoplankton) or by fragments of them (animals or macroalgae). Species taxonomically close to *G. minor* have been found associated with sea-animals^10,11^, macroalgae^12^ and even dinoflagellates^39,40^. Related to this, data of the same cruise^41^ reported highest concentrations of large phytoplankton in the deep sea, represented by living fast-sinking cells (81.5% of diatoms followed by dinoflagellates) in the Equatorial Pacific, opening the hypothesis of undiscovered microbial interactions. Interestingly, the sequencing analyses shows an absence of *G. minor* in a large surface area of the equatorial Atlantic Ocean, from 14° N to 24° S (stations 17 to 26) so, the lower presence of this species in Atlantic bathypelagic waters could be explained by its absence at the surface, as the snowfall of particles at the equator is similar between the two oceans^42^.

Despite several hints pointing to the fact that fungi arrive to the deep ocean attached to particles, we have only been able to observe this phenomenon once (Fig. 2e, f, which shows a large particle colonized by rounded cells), while most observed rounded cells appear to be free-living. There are possible methodological reasons to explain the lack of particles observed in our TSA-FISH filters (low volume, high filtering pressure); nevertheless, even in an environment full of particles, yeasts are expected to look for new resources once the ones of their transport-particle are exhausted. We propose that, although the manipulation of the samples could separate some cells from particles, part of the community live temporarily in a free-living state as evidenced by past results of flow cytometry^16^, sequencing of the 0.2-0.8 µm size fraction (often used as the free-living size fraction for bacteria, supplementary table 1) and the TSA-FISH itself.

The biomass of *G. minor* yeast cells correlates very well with the biomass of free-living bacteria, although fungal biomass was 3 orders of magnitude lower than the bacterial one. This contrasts Bochadansky et al. (2017)^9^, who found similar values for the two groups. The prokaryotic free-living pool represents the endemic part of the community and, in contrast with the particle attached assemblage, does not correlate with surface biotic variables but with bathypelagic environmental conditions^38^. This leads us to propose that the good correlation between the biomass of fungi and free-living bacteria is due either to a parallel response to the bathypelagic conditions and resources or to the utilization by fungi of carbon processed by bacteria. In fact, both cases point to an active free-living fungal community. In culture conditions, *G. minor* does not grow without some vitamins^36^, and in particular, it needs thiamine to thrive. It is possible that the prokaryotic community could be the source of these vitamins and a reliable scenario is that vitamin availability coupled with recalcitrant carbon (C1 peak) created optimal conditions for *G. minor,* which could grow and reproduce in marine deep waters.

The idea that once passively transported to the Bathypelagic layer of the Pacific, *G. minor* is capable of establishing an actively thriving population is supported by several hints. First, we know from experimental evidence in *Saccharomyces cerevisiae* that fungi can alter their membrane composition to tolerate high hydrostatic pressures^43^, which could make possible a rapid colonization of deep-sea habitats by surface strains^44^. For *G. minor* this colonization could be supported by a great availability of resources, the pool of recalcitrant carbon. Second, considering the entire dataset, not only Basidiomycota but also Ascomycota are more abundant in the aged waters. Moreover, Ascomycota peak is displaced spatially forward compared to the Basidiomycota peak, pointing to a different target substrate and reinforcing the hypothesis of an active fungal deep community. Third and more importantly, our analysis showed a strong and significant correlation between *G. minor* biomass and recalcitrant carbon (C1 peak of FDOM) in Pacific waters suggesting a possible consumption of the recalcitrant pool by *G. minor* (Fig. 6); the same relation is not significant for bacterial biomass. This possibility is in accordance with Clipson et al. (2006)^45^ who state that fungi are often better than bacteria at breaking down recalcitrant organic material. Although the biomass of *G. minor* is much lower than the biomass of bacteria, it is still possible that they both contribute to the degradation in a mutualistic rather than antagonistic way^9,46^.

*G. minor* also relates significantly to labile carbon (C3 peak of FDOM). In fact, the slope of the correlation line between the labile C3 peak and *G. minor* is steeper than in the C1 relationship in Pacific waters, which could point to faster utilization of labile than recalcitrant carbon by fungi. It has been proposed that labile matter could act as a primer for the digestion of more refractory components in a dynamic known as *priming*^7,47–49^ . Although this topic is still controversial for the aquatic environment^50^, our results could suggest a parallel consumption of Humic-like and Labile-like matter operated by *G. minor*.

The presence of the cultured plant-parasite *G. minor* has been highlighted in bathypelagic waters, especially in the Equatorial Pacific where its biomass peaks, both by species-specific TSA-FISH probe and by high-throughput sequencing. Our data suggests that in these waters *G. minor* feeds osmotrophically on the recalcitrant carbon pool, a resource with which it correlates better than free-living bacteria. Our results show, in accordance to previous works^38,51^, that the Bathypelagic Ocean is not an isolated environment but it is in continue connection with surface waters and with the land. In this regard the fungus *G. minor* shows an enormous potential to be an important active part of the global carbon cycle, pointing at the same time to the importance and necessity of more studies both on the Bathypelagic Ocean and on its fungal diversity.

## Acknowledgements

This project was supported by the Spanish Ministry of Economy and Competitiveness through projects Consolider-Ingenio Malaspina-2010 (CSD2008–00077), ALLFLAGS (CTM2016-75083-R, MINECO) and MINIME (PID2019-105775RB-I00, AEI, Spain).

We thank our fellow scientists, the crew of the R/V BIO-Hesperides and chief scientists of the different cruise legs for collaboration, a special thank goes to J.M. Gasol and X.A.G. Moran for sharing the bacterial abundance data.

## Data Availability

The sequencing dataset analysed in this present work are all belonging to Malaspina expedition and they are already published and publicly available at the European Nucleotide Archive (ENA). Amplicon sequences from 454-pyrosequencing of the deep ocean are available at www.ebi.ac.uk/ena/data/view/PRJEB9943; Illumina sequences from vertical profiles are available at www.ebi.ac.uk/ena/data/view/PRJEB23771; Illumina sequences from surface samples are available at www.ebi.ac.uk/ena/PRJEB23913 and Illumina sequences from the deep ocean are available at www.ebi.ac.uk/ena/data/view/PRJEB45014.

## Authors contribution

MCP, RM, and RL ideate the paper, IF developed the protocol and counted the samples at the microscope, MCP developed the figures and wrote the first draft. MCP, RM, RL and IF edited the final version of the paper.

