## supplementary table 1 for "A fungi hotspot deep down the ocean: explaining the presence of *Gjaerumia minor* in Equatorial Pacific bathypelagic waters"

| Station | Depth | *G. minor* % | Size fraction |
| --- | --- | --- | --- |
| 10 | 3944 | 0.00 | 0.2-0.8 |
| 23 | 3942 | 0.02 | 0.2-0.8 |
| 26 | 3847 | 0.03 | 0.2-0.8 |
| 32 | 3200 | 0.29 | 0.2-0.8 |
| 35 | 3650 | 0.15 | 0.2-0.8 |
| 41 | 4000 | 0.21 | 0.2-0.8 |
| 67 | 4000 | 0.15 | 0.2-0.8 |
| 74 | 4000 | 11.72 | 0.2-0.8 |
| 77 | 4000 | 0.17 | 0.2-0.8 |
| 82 | 2150 | 0.72 | 0.2-0.8 |
| 97 | 4000 | 1.26 | 0.2-0.8 |
| 103 | 4000 | 17.51 | 0.2-0.8 |
| 109 | 4000 | 0.99 | 0.2-0.8 |
| 112 | 4000 | 9.40 | 0.2-0.8 |
| 118 | 3200 | 1.56 | 0.2-0.8 |
| 121 | 3000 | 0.19 | 0.2-0.8 |
| 131 | 4000 | 0.90 | 0.2-0.8 |

**Table S1:** This table shows the percentage of reads belonging to ASV-363 in the size fraction 0.2-0.8 µm. This size fraction is not our primary target and for this reason these data were not included in any figure. Nevertheless, we noticed that several cells have a diameter around 0.9 µm and could be they trespass the 0.8 filter due to the filtration pressure. The presence of these cells could be an hint of a free-living life-style.
